# Immunoprecipitation of RNA-DNA hybrid interacting proteins in *Trypanosoma brucei* reveals conserved and novel activities, including in host immune evasion by antigenic variation

**DOI:** 10.1101/2023.05.11.540366

**Authors:** Mark J. Girasol, Emma M. Briggs, Catarina A. Marques, José M. Batista, Dario Beraldi, Richard Burchmore, Leandro Lemgruber, Richard McCulloch

## Abstract

RNA-DNA hybrids are widespread epigenetic features of genomes that provide a growing range of activities in transcription, chromatin and DNA replication and repair. Understanding of these diverse functions has been advanced by characterising the proteins that interact with the hybrids, with all such studies revealing hundreds of potential interactors. However, all interaction analyses to date have focused on mammalian cells, and so it is unclear if a similar spectrum of RNA-DNA hybrid interactors is found in other eukaryotes, thus limiting our understanding of the conserved and lineage-specific activities linked to these genetic structures. The African trypanosome is a compelling organism in which to address these questions. As a divergent single-cell eukaryotic parasite of the Discoba grouping, *Trypanosoma brucei* displays substantial divergence in several aspects of core biology from its mammalian host and, unusually for a protist, has well-developed tools for molecular genetic analysis. For these reasons, we used DNA-RNA hybrid immunoprecipitation coupled with mass spectrometry to reveal 602 putative interactors in *T. brucei* mammal- or insect vector-infective stage cells. We show that the approach selects for a subset of the parasite proteome and reveals a range of predicted RNA-DNA hybrid associated activities, some overlapping with similar studies in mammals. We demonstrate that loss of three factors, two putative helicases and a RAD51 paralogue, impact on *T. brucei* nuclear RNA-DNA hybrid and DNA damage levels. Moreover, loss of each affects the operation of the crucial parasite immune survival mechanism of antigenic variation. Thus, our work reveals the broad range of activities contributed by RNA-DNA hybrids to *T. brucei* biology, including new functions in host immune evasion as well as many conserved with mammals, and so likely fundamental to eukaryotic genome function.

## Introduction

RNA-DNA hybrids are ubiquitous features of genomes in all domains of life. R-loops are a form of RNA-DNA hybrid in which an RNA molecule base-pairs with one strand of double-stranded DNA, causing displacement of a DNA single-strand. Though initially found to form during transcription (Aguilera and Garcia-Muse, 2012), R-loops are increasingly known to be widespread in genomes and to have wide impacts on genome function, both positive and negative (Brickner et al., 2022; Niehrs and Luke, 2020; Petermann et al., 2022). Such activities include initiation and arrest of DNA replication (Lombrana et al., 2015; Stuckey et al., 2015; Tran et al., 2017), transcription activation and termination (Santos-Pereira and Aguilera, 2015), chromatin formation (Chedin, 2016; Zhang et al., 2017), and telomere function (Nanavaty et al., 2017a; Rippe and Luke, 2015). R-loops can result in genome instability (Aguilera and Garcia-Muse, 2012; Sollier and Cimprich, 2015; Stirling and Hieter, 2016), at least in part by generating DNA breaks, which can be harmful (Garcia-Muse and Aguilera, 2019), but can also be used functionally, such as during class switch recombination in mammalian B cells (Daniels and Lieber, 1995; Matthews et al., 2014; Yu et al., 2003). RNA-DNA hybrids and R-loops can also contribute to DNA damage repair, including of DNA double-strand breaks (Brickner et al., 2022; Ketley and Gullerova, 2020; Marnef and Legube, 2021).

A number of studies have recently begun to dissect the various activities provided by RNA-DNA hybrids and R-loops by characterising the proteins with which they interact. The earliest of these studies searched for RNA-DNA hybrid interactors in HeLa cells using immunoprecipitation with the S9.6 monoclonal antibody (Boguslawski et al., 1986) and mass spectrometry (Cristini et al., 2018), revealing hundreds of potential activities. A similar approach in mouse embryonic stem cells also revealed hundreds of putative RNA-DNA hybrid interactors (Wu et al., 2021). A distinct approach used two large, synthetic RNA-DNA hybrids and recovered >1000 proteins each from lysates of human B cells (Wang et al., 2018). Two more recent studies relied on proximity-labelling based on the DNA binding domain of RNase H1, identifying ∼300-400 proteins in immortalised human cells (Mosler et al., 2021; Yan et al., 2022). Finally, based on all these datasets, Kumer et al (Kumar et al., 2022) searched for common features of the recovered proteins and used machine learning to predict RNA-DNA hybrid interacting proteins across the human proteome. Together, these studies have revealed a wealth of potential RNA-DNA hybrid and R-loop associated activities. However, all these studies are limited to mammals, and no study has asked if similar or distinct activities are found in other eukaryotes. Here, we have adapted the DNA-RNA immunoprecipitation-mass spectrometry (DRIP-MS) approach of Cristini et al (Cristini et al., 2018) to explore the RNA-DNA hybrid interactome of the protozoan parasite, *Trypanosoma brucei*, where mapping of R-loops has predicted both conserved and diverged genomic activities (Briggs et al., 2018b).

The genome of *T. brucei* is unusual for a eukaryote in several respects (Berriman et al., 2005; Muller et al., 2018). All but two of the ∼8000 protein-coding genes in *T. brucei*’s genome ‘core’ (see below) are transcribed by RNA Polymerase (Pol) II from multigene clusters, each of which contains potentially hundreds of genes and is transcribed from a single, still only partly understood transcription start site (Cordon-Obras et al., 2022; Siegel et al., 2009; Staneva et al., 2022; Staneva et al., 2021; Wedel et al., 2017). Unlike in bacteria, genes in such operon-like transcription units appear not be functionally related (Daniels et al., 2010). This arrangement means that the genome core contains relatively limited content that is not traversed by RNA Pol-II, and mature mRNAs are generated from pre-mRNA transcripts by extensive, coupled trans-splicing and polyadenylation (Siegel et al., 2011). In addition, each multigene transcription unit has a single transcription termination site, which contains a novel base, termed J, that acts to recruit a number of termination factors (Kieft et al., 2020). This unusual organisation of gene expression appears to also reflect DNA replication organisation, since mapping sites of replication initiation, termed origins, reveals close overlap with transcription start and termination sites (Devlin et al., 2016; Tiengwe et al., 2012). Furthermore, RNAi against a subunit of the origin recognition complex, which defines origins (Marques et al., 2016), suggests functional interaction between the replication and transcription machineries (Tiengwe et al., 2012). All the above aspects of gene expression appear conserved with the wider grouping of kinetoplastids, while other aspects of the *T. brucei* genome may be specific. Survival of African trypanosomes in the mammal relies on a process termed antigenic variation, which involves continuous changes in the identity of the surface expressed ‘coat’, which is composed of a single Variant Surface Glycoprotein (VSG) in a single cell at a given time (Faria et al., 2022; McCulloch et al., 2017). Switching from one VSG coat to another in the mammal relies on both transcriptional changes between approximately 15 telomeric *VSG* transcription sites, termed bloodstream expression sites (BES), and recombination reactions that move silent *VSG* genes into the actively transcribed BES (Faria et al., 2022; McCulloch et al., 2015). Each BES is also a multigene transcription unit but is, remarkably, transcribed by RNA Pol-I from a promoter that shares some homology with those at rRNA gene clusters (Gunzl et al., 2003). Recombination relies on a huge archive of 1000s of silent *VSG*s, which are mainly found in arrays that occupy the chromosome subtelomeres (Berriman et al., 2005; Cross et al., 2014; Marcello and Barry, 2007). Each chromosome in *T. brucei* thus comprises a highly transcribed core and predominantly untranscribed subtelomeres, with chromosome conformation capture and ATAC-seq analyses indicating that the two genome compartments rarely interact and display differing levels of chromatin-mediated compaction (Muller et al., 2018).

Previous work has mapped R-loops in *T. brucei*, revealing that their localisation and potential functions span all the above aspects of the genome. DNA-RNA immunoprecipitation and sequencing (DRIP-seq) indicates that R-loops localise to the start and, to a lesser extent, end of the RNA Pol-II multigene clusters, as well as to intra-cluster regions of splicing and polyadenylation (Briggs et al., 2018b). The same approach showed that R-loops localise to the single centromere found in each chromosome. Examining the impact of loss of *T. brucei* RNase H1 or the A subunit of RNase H2 (both of which are RNA-DNA endonucleases that remove RNA from RNA-DNA hybrids) (Cerritelli and Crouch, 2009) revealed DNA damage associated with RNA Pol-II transcription initiation and R-loop accumulation and DNA damage within the *VSG* BESs, which causes loss of VSG expression control and increased *VSG* switching (Briggs et al., 2019; Briggs et al., 2018a; Eisenhuth et al., 2021). Loss or overexpression of RNase H1 or RNase H2 also alters levels of telomeric RNA-DNA hybrids and affects *VSG* expression and switching (Briggs et al., 2019; Briggs et al., 2018a; Nanavaty et al., 2017b; Saha et al., 2021). To begin to explore this diverse range of RNA-DNA hybrid functions in *T. brucei*, we show here that DRIP-MS recovers a similarly large number of interacting proteins as is observed in mammalian cells. Amongst these putative interactors we can identify functions conserved in mammals, including ribosome- and mRNA-associated factors and helicases, as well as activities that may reflect the unusual *T. brucei* genome, including histone variants and centromere-binding kinetochore proteins. We provide functional analysis of three DRIP-MS factors and show that loss of any of them increases nuclear damage and RNA-DNA hybrid levels, as well as altering VSG expression. One of these factors is a RAD51-related protein previously described to act in *VSG* recombination (Dobson et al., 2011; Proudfoot and McCulloch, 2005), while the two others are putative helicases that have never been functionally examined in *T. brucei*.

## Results and discussion

### Identifying the *T. brucei* RNA-DNA hybrid interactome

RNA-DNA hybrids and associated proteins were enriched through immunoprecipitation (DRIP) using the S9.6 antibody, which recognizes RNA-DNA hybrids at an affinity as low as 0.6 nM (Phillips et al., 2013). Since S9.6 can also recognise double-stranded RNA (Hartono et al., 2018), nuclei were first released from the cells by homogenization after mild lysis. The released nuclei were then sonicated to fragment the genomic DNA and limit co-precipitation of non-specific proteins during RNA-DNA hybrid recovery. DRIP was then performed on native chromatin, without cross-linking, in the presence of RNase A to deplete nuclear RNA not associated with DNA, and thereby limit DRIP of RNA binding proteins. In addition, half of the nuclear extracts were treated with benzonase prior to DRIP. Benzonase digests all forms of nucleic acids (Moreno et al., 1991), including RNA/DNA hybrids, and so these DRIP samples provided controls for recovery of non-specific proteins by S9.6. DRIPs, plus benzonase controls, were performed for both *T. brucei* procyclic forms (infective to the insect vector; two biological replicates) and bloodstream forms (mammal-infective; four replicates) grown in culture. Fig. S1A shows the results of such DRIPs after gel separation, revealing that treatment with benzonase caused loss of many detectable DRIP proteins from both PCF and BSF cells. To characterize the recovered proteins, matched sections of gels for both benzonase-treated and non-treated DRIP samples from all independent replicates were analysed by mass spectrometry (MS).

To identify putative RNA-DNA hybrid interacting proteins from the label-free DRIP-MS data, emPAI values of all proteins were compared, in each DRIP sample, to protein emPAI values in their cognate benzonase-treated control DRIP samples and log2 enrichment determined. In total, 602 proteins were enriched in at least one DRIP-MS replicate relative to the benzonase controls: 463 and 351 putative RNA-DNA hybrid-interacting proteins in BSF and PCF cells, respectively (Fig. 1, Table S1). To ask if the DRIP was selective, fold BSF/PCF enrichment of the 602 DRIP-MS enriched proteins was compared with whole-cell proteomic data from Butter et al. (Butter et al., 2013) (Fig. S1B). No correlation in BSF/PCF enrichment ratios was seen between these two studies, indicating DRIP enriched for a non-random set of *T. brucei* proteins. Cellular compartment gene ontology (GO) term enrichment analysis was next performed on the putative RNA-DNA hybrid interactors. Consistent with nuclear enrichment prior to DRIP, and with the likelihood that most RNA-DNA hybrids will form on the nuclear genome, predicted nuclear and nucleolar proteins showed significant GO term enrichment in the PCF data, and nucleolar proteins were significantly enriched in the BSF data (Fig. 1A). In contrast, nuclear envelope and plasma membrane proteins were under-represented. Though cytoplasmic proteins were not enriched (Fig. 1A), they represented the majority of proteins recovered from each life cycle stage, perhaps suggesting recovery of proteins that are not RNA-DNA hybrid interactors; for instance, ribosomal proteins were prominent in the DRIP-MS dataset (Table S1), possibly due to S9.6 recognition of double-stranded RNA. However, the recovery of ribosomal proteins may also reflect growing evidence for roles of R-loops in rRNA transcription and processing, leading to ribosome biogenesis (Abraham et al., 2020; Lawrimore et al., 2021), and is consistent with machine-learning predictions of human RNA-DNA hybrid interacting protein types (Kumar et al., 2022). Moreover, recent work has shown R-loops formed in the nucleus can be found in the cytoplasm of human cells (Crossley et al., 2023), where it is possible that they interact with cytoplasmic proteins. Indeed, below we provide evidence that at least one *T. brucei* RNA-DNA hybrid interactor predominantly localises to the cytoplasm but can provide nuclear activities.

**Figure 1.**
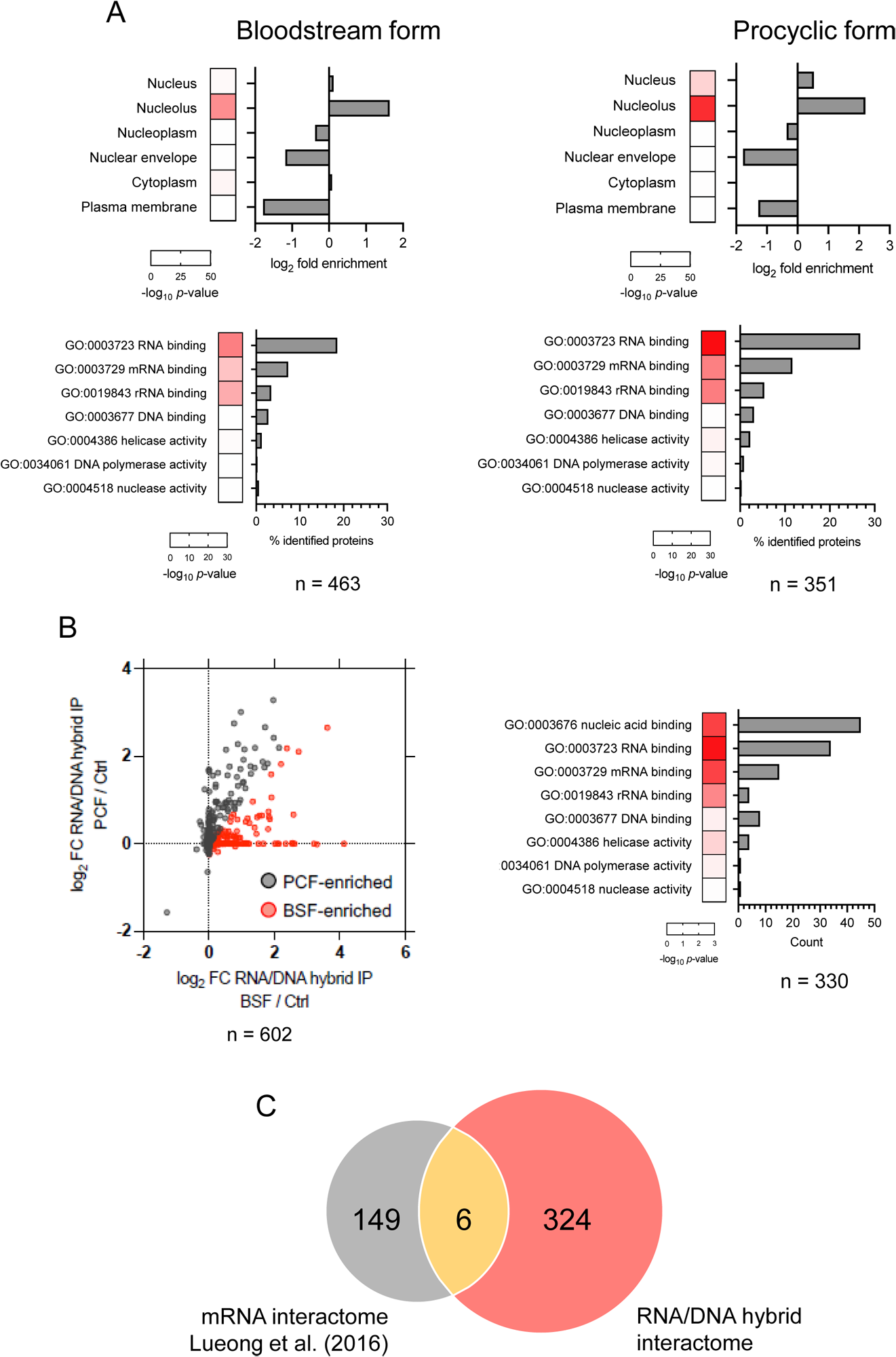
Characterisation of the *T. brucei* RNA-DNA hybrid interactome. (A) The top two panels show cellular compartment GO term analysis of proteins recovered by RNA-DNA hybrid immunoprecipitation and mass spectrometry (DRIP-MS) from bloodstream (BSF) and procyclic form cells (PCF), while the lower panels show molecular function GO term analysis of the same datasets. For cellular compartment analysis, fold enrichment relative to the proteome is indicated, while for molecular function analysis the different categories are shown as percentage of total recovered proteins (n); for both, Bonferroni-adjusted p-values are shown as heatmaps. (B) Scatter plot of log_2_-transformed mean emPAI values of BSF and PCF DRIP-MS proteins relative to benzonase controls, with those proteins enriched in BSF cells shown in red and those enriched in PCF cells shown in grey. (C) Molecular function GO term analysis of BSF-enriched proteins (details as in A). (D) Comparison of the BSF RNA/DNA hybrid interactome and mRNA interactome obtained from a study by Lueong et al. (2016a).

### Conserved eukaryotic RNA-DNA hybrid-associated functions predicted by DRIP-MS

Next, molecular function GO term analysis was used to predict activities of the proteins recovered by DRIP (Fig. 1A). Multiple proteins with predicted nucleic acid–binding activities were enriched, mirroring previous interactome data from mammalian cells (Cristini et al., 2018) and consistent with selective S9.6 immunoprecipitation. RNA-binding proteins showed the most significant enrichment in both the BSF and PCF DRIP interactomes. Pronounced enrichment of mRNA-binding proteins is a common feature of our data and that of mammalian HeLa cells (Cristini et al., 2018), and here might reflect the ubiquitous localisation of R-loops with pre-mRNA processing regions in RNA Pol-II polycistrons (Briggs et al., 2018b). Recovery of mRNA binding proteins was non-random (see below) and may suggest the hybrids provide activities linked to mRNA maturation, such as provided by the numerous predicted helicases recovered (Table S1). rRNA-binding proteins were also enriched, as also seen by Wu et al in mouse cells (Wu et al., 2021), providing further evidence of ribosome-associated R-loop functions (Fig.1A).

A number of classes of recovered factors provide evidence for RNA-DNA hybrids acting during gene expression in *T. brucei*. All four known *T. brucei* histone variants were enriched in the DRIP-MS data (Table S1): H2A.Z and H2B.V, which localise at transcription initiation regions, and H4.V and H3.V, which localise to termination regions (Siegel et al., 2009). Thus, the DRIP-MS data reinforces previous *T. brucei* DRIP-seq mapping that showed enrichment of RNA-DNA hybrids at transcription start sites and, to a lesser extent, termination sites, indicating R-loop roles in gene expression organisation (Briggs et al., 2018b). In mammalian cells, RNA-DNA hybrids were found to interact with histone H3 (Cristini et al., 2018), and the hybrids are known to recruit histone H3 modifications at promotor regions (Sanz et al., 2016). R-loops have not been mapped in further trypanosomatids but such roles may be conserved, as the histone variants show the same localisations (Anderson et al., 2013; Roson et al., 2022). How the deposition and functions of trypanosome R-loops and histone variants intersect is unknown, and the potential contribution of the hybrids to transcription remains unclear. However, although the DRIP did not recover RNase H1 or RNase H2 (in common with Crisitini et al (Cristini et al., 2018)), loss of RNase H2A causes pronounced DNA damage accumulation at transcription start sites (Briggs et al., 2019). In addition, immunoprecipitation of the histone methyltransferases DOT1A or DOT1B reveals interaction with RNase H2 (Eisenhuth et al., 2021; Staneva et al., 2021), with DOT1A-RNase H2 activity potentially resolving R-loops at transcription termination sites containing RNA Pol-III genes (Staneva et al., 2021). We did not detect either DOT1A or DOT1B in the DRIP-MS data, perhaps suggesting the methyltransferases do not directly interact with RNA-DNA hybrids.

Adding to the above gene expression-associated activities, two of four *T. brucei* (Mani et al., 2011) Alba proteins were recovered: ALBA3 and ALBA4 (Table S1). Alba proteins are found in both archaea and eukaryotes and bind DNA and/or RNA (Goyal et al., 2016). Studies to date have shown *T. brucei* ALBAs to be cytoplasmic and possess RNA binding roles in translation (Bevkal et al., 2021; Melo do Nascimento et al., 2021; Subota et al., 2011), and so interaction with RNA-DNA hybrids may be surprising. However, in *Arabidopsis thaliana* ALBA1 binds RNA-DNA hybrids and ALBA2 interacts with the displaced single-stranded DNA, and together acts as an R-loop reader complex (Yuan et al., 2019). Additionally, the four ALBA proteins found in *Plasmodium falciparum* bind both DNA and RNA (Chene et al., 2012), and so the prediction of RNA-DNA hybrid interaction for *T. brucei* ALBA proteins may indicate previously unknown activities.

DNA-binding proteins showed only weak evidence for enrichment (Fig. 1A), unlike the pronounced enrichment seen in mammal DRIP-MS data (Cristini et al., 2018). However, this grouping included some notable factors when specific functions were examined, suggesting roles for *T. brucei* RNA-DNA hybrid interactors in chromosome functions. Previously, we described enrichment of R-loops at *T. brucei* centromeres using DRIP-seq (Briggs et al., 2018b). Consistent with such localisation, three kinetochore proteins (Akiyoshi and Gull, 2014) were recovered by DRIP: KKT1, KKT3, and KKT19 (Table S1). Though R-loops have also been shown to localise to centromeres in yeast (Mishra et al., 2021), mammals (Kabeche et al., 2018) and plants (Liu et al., 2023; Liu et al., 2021), it is unclear how they might contribute to, or indeed impede, centromere function. Indeed, most trypanosomatid kinetochore subunits show no evidence of orthology with kinetochores in other eukaryotes (Akiyoshi and Gull, 2014; D’Archivio and Wickstead, 2016). Intriguingly, *T. brucei* KKT3 is thought to be one of two kinetochore proteins that are positioned most proximal to the centromere (D’Archivio and Wickstead, 2016), where they localise throughout the cell cycle (Akiyoshi and Gull, 2014). In addition, they contain novel domains for centromere association (Marciano et al., 2021) and recruit KKT1 to assemble the kinetochore (Ishii et al., 2022). DRIP recovery of KKT3 and KKT1 may then suggest that *T. brucei* centromeric R-loops help maintain and guide interaction of the kinetochore to the centromere throughout the cell cycle. Alternatively, R-loops may provide epigenetic definition of the *T. brucei* centromere in the absence of the histone H3 variant, CENP-A (Akiyoshi and Gull, 2013). SMC (structural maintenance of chromosome) proteins are ATPases found in all domains of life and are core subunits of larger protein complexes needed to organise the genome through conformational change (Hassler et al., 2018). Two of these complexes in eukaryotes are cohesin, which contains Smc1 and Smc3, and condensin, containing Smc2 and Smc4. Here, DRIP recovered both *T. brucei* SMC2 and SMC4 (Table S1). Unlike for cohesin (Gluenz et al., 2008; Landeira et al., 2009), no work to date has described condensin functions in any kinetoplastid, and so the prediction of R-loop interaction may provide a route to examine where and how the complex acts in *T. brucei*, where chromosome condensation during mitosis appears to be minimal (Yamin et al., 2022).

Beyond the above protein cohorts, DRIP-MS implicated a number of further, less easy to predict RNA-DNA hybrid-associated activities. For instance, several protein kinases were recovered (Table S1), including three NEK family kinases (Jones et al., 2014), which have diverse roles including in cell cycle control and DNA damage repair (Pavan et al., 2021). Though mitochondrial proteins are likely to be under-represented in our approach, several kinetoplast proteins were recovered (Table S1), including two DNA Pols (IC and Beta-PAK) (Maldonado et al., 2021; Miller et al., 2020) and a putative RNA-editing nuclease. Finally, nearly a third of the proteins (180) recovered by DRIP are annotated as hypothetical or hypothetical conserved (Table S1), and so no functions can be predicted currently.

### Searching for RNA-DNA hybrid interactor roles in *T. brucei* antigenic variation

Amongst the wealth of potential RNA-DNA hybrid interactors revealed by DRIP-MS, we decided to ask if activities associated with antigenic variation could be identified, since R-loops are involved in the pathway in ways that are not yet clear (Damasceno et al., 2021; Saha et al., 2020). In addition, we reasoned that some R-loop activities that act in antigenic variation could be unique to African trypanosomes and might therefore yield a means to impede this crucial survival mechanism (Faria et al., 2022).

Despite DRIP-MS not detecting direct RNase H2A or DOT1B interaction with RNA-DNA hybrids, there is clear evidence of functional interaction between these proteins and their involvement in antigenic variation: loss of DOT1B or RNase H2A results in similar changes in R-loop levels, DNA damage levels and Variant Surface Glycoprotein (VSG) expression alterations (Briggs et al., 2019; Eisenhuth et al., 2021). In fact, a connection between R-loops and VSG expression may also extend to the histone variants recovered by DRIP-MS, given similarities in changes to *VSG* transcription control after loss of either RNase H (Briggs et al., 2019; Briggs et al., 2018a) or H3.V and/or H4.V (Kim, 2021; Muller et al., 2018; Reynolds et al., 2016). Intriguingly, DRIP-MS data predicted VEX1 as a hybrid interactor, perhaps suggesting even more widespread roles for R-loops in VSG expression (Table S1), consistent with the similarity in de-repression of silent *VSG* expression sites after loss or overexpression of VEX1 (Glover et al., 2016), in RNase H1 mutants (Briggs et al., 2018a), and after RNase H2A RNAi (Briggs et al., 2019; Eisenhuth et al., 2021). One explanation for DRIP recovery of VEX1 may be that the protein localises within or proximal to the telomeres of *VSG* expression sites (Faria et al., 2019), where RNA-DNA hybrids are present (Briggs et al., 2019; Briggs et al., 2018a; Nanavaty et al., 2017b). We did not detect the other components of the VEX complex (Faria et al., 2019), such as VEX2, however, and so these data may suggest a specific R-loop role of VEX1 (Faria et al., 2021).

To ask if other RNA-DNA interacting proteins functionally link R-loops and antigenic variation, including by previously unknown mechanisms, we first attempted to compare the abundance of DRIP recovered proteins in BSF and PCF cells, asking which are more prevalent in the former, since VSG is not expressed in the latter (Fig. 1B). A total of 330 putative BSF-enriched interactors were seen at a very permissive selection level (log2 fold-change BSF/PCF > 0). Molecular function GO term analysis of this subset did not reveal any clear difference to all 602 proteins recovered (Fig. 1B). Nonetheless, we were now more able to compare the RNA/DNA hybrid interactome with the BSF mRNA proteome generated by Lueong et al (Lueong et al., 2016b) (Fig. 1B), since the GO term ‘mRNA binding’ was consistently enriched in all DRIP-MS analyses (Fig. 1A,B). Overlap between the two proteomic datasets was very limited, further showing that S9.6 immunoprecipitation recovers a non-random selection of *T. brucei* proteins.

Given that comparing predicted RNA-DNA hybrid interactomes between two life cycle stages did not yield any obvious difference in enrichment patterns, we decided to narrow the search based on two criteria: looking for proteins with annotations of relevant predicted activity; and DRIP-MS indication of recovery only from BSF cells. Amongst the proteins that fulfilled these criteria, four were selected for further analysis. The first two proteins were chosen because they are known to act in *T. brucei* VSG switching: RAD51 (Tb927.11.8190) (Glover et al., 2013; Jehi et al., 2014; McCulloch and Barry, 1999) and the RAD51 paralogue, RAD51-3 (Tb927.11.2550) (Dobson et al., 2011; Proudfoot and McCulloch, 2005). RAD51 from yeast and mammals has previously been shown to bind RNA-DNA hybrids (Feretzaki et al., 2020; Wahba et al., 2013), and we describe the *T. brucei* RAD51-directed connection between R-loops and VSG switching elsewhere (Girasol et al, BioRxiv). RAD51 paralogues are related to RAD51 and provide a range of activities in DNA damage repair and replication (Bhattacharya et al., 2022; Bonilla et al., 2020), but no work to date has suggested interaction with RNA-DNA hybrids. The two other proteins, encoded by Tb927.3.2600 and Tb927.3.5440 (Fig. 2), were chosen as they provide relevant predicted functions (below) that have not been experimentally examined in *T. brucei* to date.

**Figure 2.**
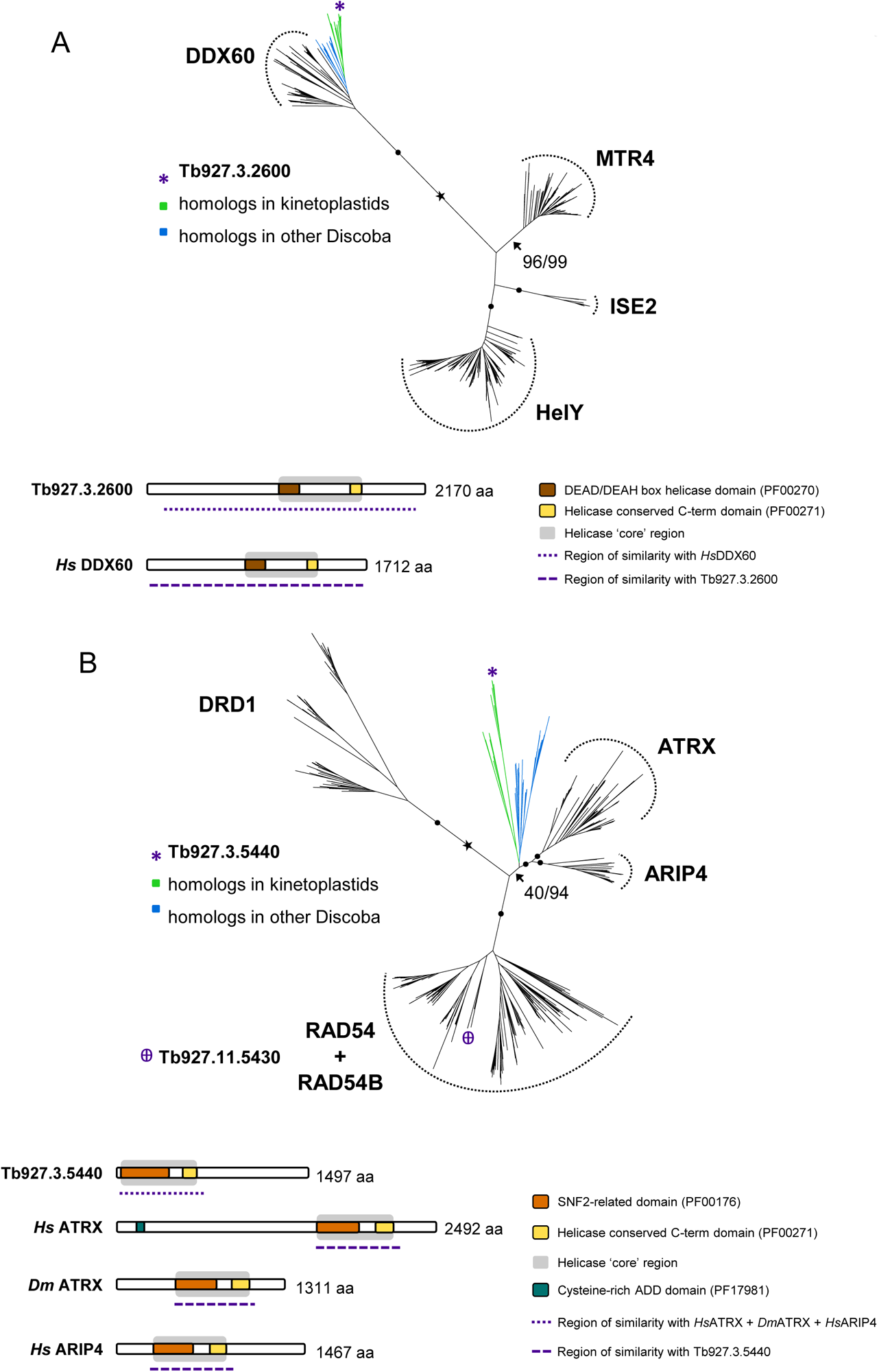
Identification of the putative RNA-DNA hybrid interactors encoded by Tb927.3.2600 and Tb927.3.5440. Phylogenetic and domain analyses are shown for Tb927.3.2600 (A) and Tb927.3.5440 (B). A maximum-likelihood phylogenetic tree of Tb927.3.2600 homologs found in Discoba and representatives of selected subfamilies within the Ski2-like family of SF2 helicases places Tb927.3.2600 within the DDX60 branch with maximal support; branch support values are SH-aLRT(%)/UFBoot2(%); filled circles represent 100/100 support and the estimated root is indicated with a star. Domain organisation of Tb927.3.2600 and its *Homo sapiens* (*Hs*) homolog DDX60 (UniProt: Q8IY21) reveals that sequence similarity between both proteins extends beyond the helicase core. A maximum-likelihood phylogenetic tree of Tb927.3.5440 homologs found in Discoba and representatives of selected subfamilies within the Snf2 family of SF2 helicases suggests that Tb927.3.5440 does not belong in one of the pictured subfamilies but is likely evolutionary closer to ATRX and ARIP4. Tb927.11.5430 (highlighted) is a *T. brucei* homolog of RAD54/RAD54B. Branch support values and estimated root are as shown for Tb927.3.2600. Domain organisation of Tb927.3.5440 and putative homologs in *H. sapiens* (ATRX, UniProt: P46100; ARIP4, UniProt: Q9Y4B4) and *Drosophila melanogaster* (ATRX, UniProt: Q9GQN5) show that sequence similarity is limited to the helicase core and short flanking regions.

Tb927.3.2600 encodes one of a number of putative ATP-dependent DExD-box RNA helicases (DDXs) recovered by DRIP (Fig. 2A, Table S1). DDXs are one grouping of helicases within a larger superfamily (SF2) (Fairman-Williams et al., 2010) and, in other eukaryotes, a number of DDXs have been shown to act on R-loops: DDX21, DDX23, DDX38B and DDX41 each limit DNA damage during transcription (Mosler et al., 2021; Perez-Calero et al., 2020; Song et al., 2017; Sridhara et al., 2017); DDX19 is a nucleopore-associated factor that can translocate to the nucleus and act with the kinase ATR during transcription-replication clashes (Hodroj et al., 2017a; Hodroj et al., 2017b); DDX1 contributes to immunoglobulin class switch recombination (Ribeiro de Almeida et al., 2018); and DDX1, DDX5 and DDX18 act on R-loops associated with DNA damage, including through interaction with DNA repair factors (Li et al., 2016; Lin et al., 2022; Saha et al., 2022; Yu et al., 2020). Many of these helicases were recovered by DRIP-MS from mouse cells (Wu et al., 2021). Homology searches and phylogenetic analyses strongly suggest the *T. brucei* Tb927.3.2600-encoded DDX to be a homologue of DDX60 (Fig. 2A) and a member of the Ski2 helicase family (Fairman-Williams et al., 2010), which has not been implicated in R-loop functions in any eukaryote and has only been functionally characterised in *T. brucei* through its putative interaction with the stress response mRNA binding factor, MKT1(Ooi et al., 2020; Singh et al., 2014). The syntenically encoded protein in *T. cruzi* has recently been shown to be part of 43S preinitiation complex of the assembling ribosome (Bochler et al., 2020), an association not described in other eukaryotes, including mammals, where DDX60 is non-essential and has been instead implicated in antiviral activities and cancer (Geng et al., 2022; Sadic et al., 2022). The novelty of *T. cruzi* DDX60 interaction with the ribosome appears to be reflected in structural features not found in its mammalian orthologues (Bochler et al., 2020).

Tb927.3.5440 has been annotated (tritrypdb.org) as encoding a putative SNF2 DNA repair protein, merely suggesting that it belongs to the large Snf2 family of helicases (Fairman-Williams et al., 2010) whose members provide a wide range of activities (Hopfner et al., 2012; Joseph et al., 2020), including chromatin remodelling, transcription and DNA repair. Homology searches and phylogenetic analyses suggest the protein encoded by Tb927.3.5440 belongs to a somewhat distinct Discoba grouping that is most closely related to ATRX, which is widely distributed in eukaryotes, and ARIP4, which appears limited to animals (Fig. 2B). No function has been ascribed to *T. brucei* ATRX or its relative, RAD54/B (encoded by Tb927.11.5430; Fig. 2B), which was not detected in the DRIP-MS data (Table S1). However, ATRX in other eukaryotes has been shown to have roles in alternative lengthening of telomeres (Bhargava et al., 2022; Geiller et al., 2022; Kim et al., 2019), in homologous recombination pathway selection (Elbakry et al., 2021; Juhasz et al., 2018), and in suppression of R-loops in transcribed telomeres (Nguyen et al., 2017; Toubiana and Selig, 2018; Yan et al., 2022). Many of such roles could be consistent with RNA-DNA hybrid functions at the intersection of transcription and recombination in telomeric BESs during VSG switching in *T. brucei*.

### Loss of RAD51-3, DDX60 or ATRX leads to increased nuclear DNA damage

To begin to test the functions of the three predicted RNA-DNA interactors, we engineered MiTat1.2 BSF *T. brucei* cells to permit genetic modification via CRISPR-Cas9 [63]. Using these cells, we generated variant parasites expressing each protein as a translational fusion with mNeonGreen (mNG) and attempted to make null mutants by replacing each allelic ORF with antibiotic resistance markers.

Both alleles of all genes were successfully tagged with mNG: RAD51-3 and ATRX at the C-terminus, and DDX60 at the N-terminus (Figs. 3 and S2). In each case PCR showed integration of *BSD* and *NEO* constructs, with concomitant loss of the wild type untagged allele, and western blotting using anti-mNG antibody showed expression of fusion proteins of the expected size (Fig. S2). Tagging of RAD51-3 or DDX60 did not impair parasite growth in culture (Fig. S2A,B), whereas mNG appeared to at least partially impede ATRX protein function, since the tagged cells exhibited a growth defect compared with parental TbCas9/T7 cells (Fig. S2C). Live fluorescence microscopy revealed nuclear localization of mNG::RAD51-3 and mNG::ATRX (Fig. 3A,C) in all cell cycle stages (Fig. S2A,C). Fluorescence signal for mNG::RAD51-3 appeared more focal in cells with 1Ne1K and 1N2K compositions of nucleus (N) and kinetoplast (K) staining (see Figs. 3 and S2 for explanation), perhaps indicating recruitment to subnuclear loci during DNA replication (Fig. S2A). In all cell cycle stages DDX60::mNG signal was cytoplasmic (Figs. 3B and S2B), perhaps consistent with a ribosomal function (Bochler et al., 2020). Localisation of each protein in BSF cells essentially matches what is seen in PCF cells (Billington et al., 2023), suggesting conserved roles in at least these two life cycle stages.

**Figure 3.**
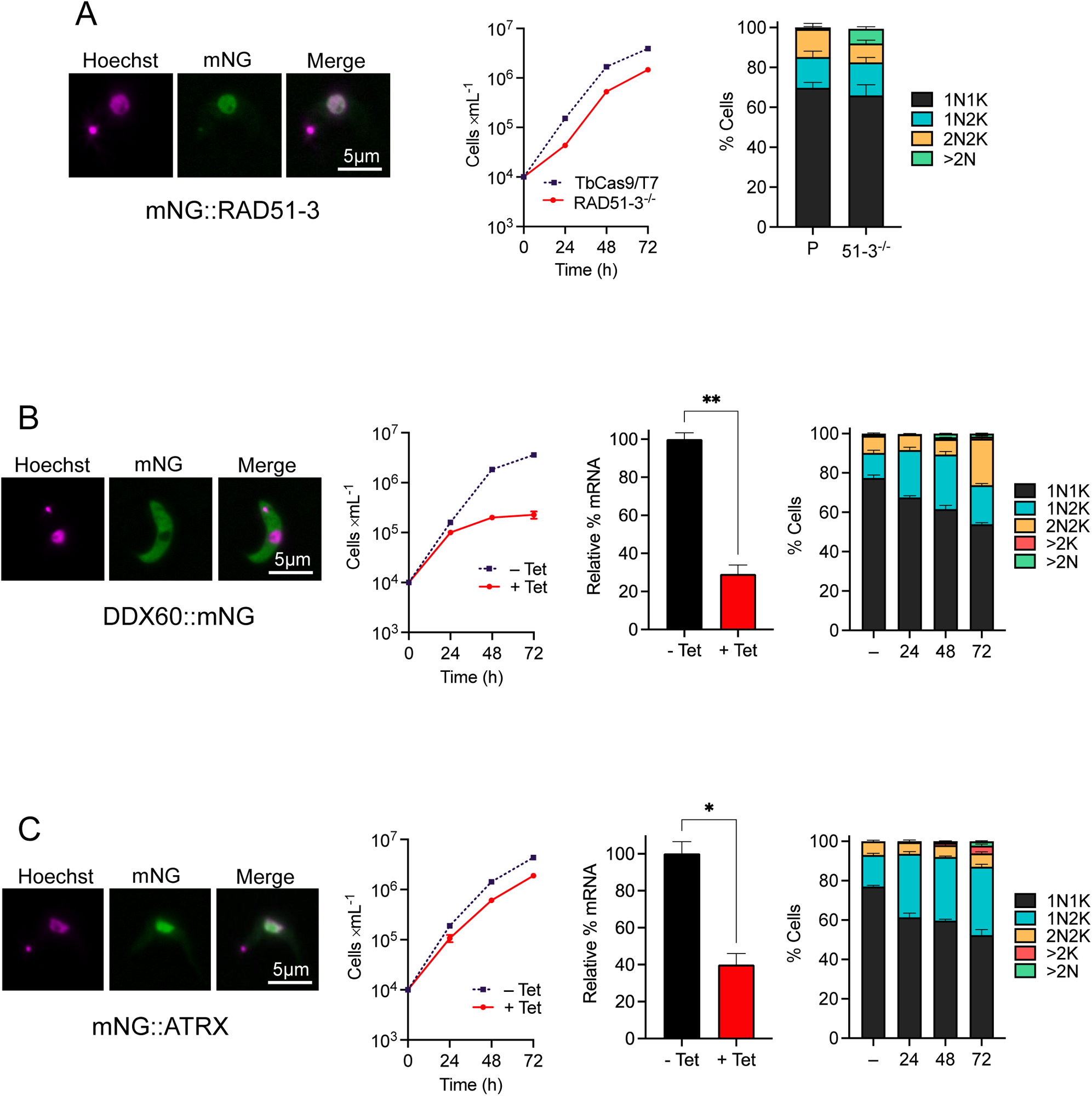
Functional characterisation of *T. brucei* RAD51-3, ATRX and DDX60. For each of RAD51-3 (A), DDX60 (B) and ATRX (C), the following are shown (left to right): representative microscopy images of live fluorescence imaging of a cell expressing the proteins as fusions with mNeonGreen (mNG; scale bar = 5 µm); growth after loss of expression (for RAD51-3 this is a comparison of parental TbCas9T7 cells relative to null mutants (-/-), while for DDX60 and ATRX RNAi induced (+ Tet) and uninduced (-Tet) cells are shown, including relative RNA levels after 24 hours of RNAi induction (uninduced RNA level was set at 100%); cell cycle profile of parental and -/- cells (RAD51-3), or before and after RNAi induction for 24, 48, and 72 hours (DDX60 and ATRX), as determined through DAPI staining of nucleus (N) and kinetoplast (K). For growth analysis, error bars represent SEM from three independent experiments. For RT-qPCR to determine RNA levels, error bars show SEM from two independent experiments and statistical significance was determined using t-test (*p<0.05). For cell cycle analysis, values are shown as a proportion of cells with specific N-K configurations (1N1K, 1N2K, 2N2K, >2K, others, e.g., >2N) in a cell population (>300 cells); error bars represent SEM from three independent experiments.

A *RAD51-3* null mutant (*RAD51-3*-/-) was generated in a single round of transfection, with PCR demonstrating replacement of both WT alleles with *BSD* and *NEO*, and RT-PCR showing loss of *RAD51-3* transcript (Fig. S3A). CRISPR-mediated deletion of *RAD51-3* confirms previous observations that the paralogue is not essential in *T. brucei* (Proudfoot and McCulloch, 2005), which may differ from *Leishmania* (Damasceno et al., 2020; Genois et al., 2015). The absence of RAD51-3 did result in a growth defect, however (Fig. 2A), which may be explained by the *RAD51-3*-/- mutants showing an accumulation of cells with more than two nuclei (>2N; 7.4±1.0% in -/-, 0.8±0.5% in parental) and a reduction in the proportion of 2N2K cells (9.4±1.7% in -/-, 14±2.9% in parental), suggesting a mitotic defect. Attempts to make null mutants of *ATRX* or *DDX60* failed. For *DDX60*, double antibiotic-resistance transformant clones were recovered, but all retained an intact ORF and displayed improper integration of *BSD* (Fig. S3B). Attempts to remove even just a single *ATRX* allele failed to yield viable antibiotic-resistant clones. To examine functions, we instead used tetracycline-inducible RNAi (Alsford and Horn, 2008). For both genes a growth defect emerged from 24 h after RNAi induction, though this was more pronounced for *DDX60* than *ATRX* (Fig. 3B,C). Concomitant with the growth defects emerging, DAPI staining revealed perturbation in the DNA content of cells within the populations. For *ATRX*, the most pronounced change was an increase in 1N2K cells (32% in induced, 16% in uninduced at 24 h; Fig. 3C), with an associated reduction in 1N1K cells, suggesting a stall in S/G2 phase. After 72 h some cells harbouring more than two kinetoplasts could be seen (4% in induced, 0% in uninduced), indicating kDNA replication and division can occur to at least some extent. The cell cycle defect after *DDX60* RNAi was distinct (Fig. 3B), with loss of 1N1K cells associated initially with accumulation of 1N2K cells (24-48 h post-induction) and later by an increase in the proportion of 2N2K cells (72 h). These effects may be explained by death of S/G2-stalled cells from 24-48 hrs, as there was little increase in cell density at these time points (Fig. 3B), or by an S/G2 stall caused by DDX60 loss that is not absolute, with some cells progressing into but not through mitosis.

To ask if the loss of the three putative RNA-DNA hybrid interactors affects nuclear genome integrity, we tested for levels of Thr130-phoshorylated histone H2A (γH2A), which is a marker for nuclear DNA damage (Glover and Horn, 2012). Western blots indicated an increased level of yH2A in *RAD51-3-/-* cells compared with parental, while yH2A levels increased after 72 h of RNAi against *DDX60* or *ATRX* (Fig. 4A). To explore these effects further, γH2A was localised and quantified by immunofluorescence (Fig. 4B). An increase in the proportion of cells with γH2A-positive nuclei was detected in *RAD51-3*-/- cells (∼42% compared with ∼5% in parental). Moreover, yH2A nuclear signal in the mutants was notably focal (Fig. 4B), suggesting discrete DNA damage accumulation and perhaps reflecting the localisation of mNG::RAD51-3 protein (Fig. S2A). Whether or not these effects are related to replication-associated DNA damage observed after loss of RAD51-3 in *L. major* is unclear (Damasceno et al., 2020). Accumulation of nuclear yH2A signal followed the growth defects seen after RNAi of *DDX*60 or *ATRX* (Fig. 4B, Fig. 3B,C): for the former, no change in the proportion of cells harbouring γH2A signal was seen 24 h after RNAi induction, whereas the signal increased significantly by 48 h and remained essentially the same at 72 h (22% and 25%, respectively); for ATRX, the proportion of cells expressing γH2A increased significantly 24 h post-induction (∼12% in induced, ∼5% in uninduced) and continued to increase from 48-72 h (∼17% to ∼34%, respectively). In both cases, yH2A signal was distinct from that seen in *RAD51-3-/-* cells, in that it appeared throughout the nucleus (Fig. 4B). Nonetheless, loss of either of these factors also resulted in nuclear DNA damage, which is perhaps most surprising for DDX60, as localisation of DDX60::mNG suggested it is cytoplasmic (Figs. 3 and S2). Whether these data indicate an undetected population of nuclear DDX60, or if the protein can dynamically move between the nucleus and cytoplasm, is unclear.

**Figure 4.**
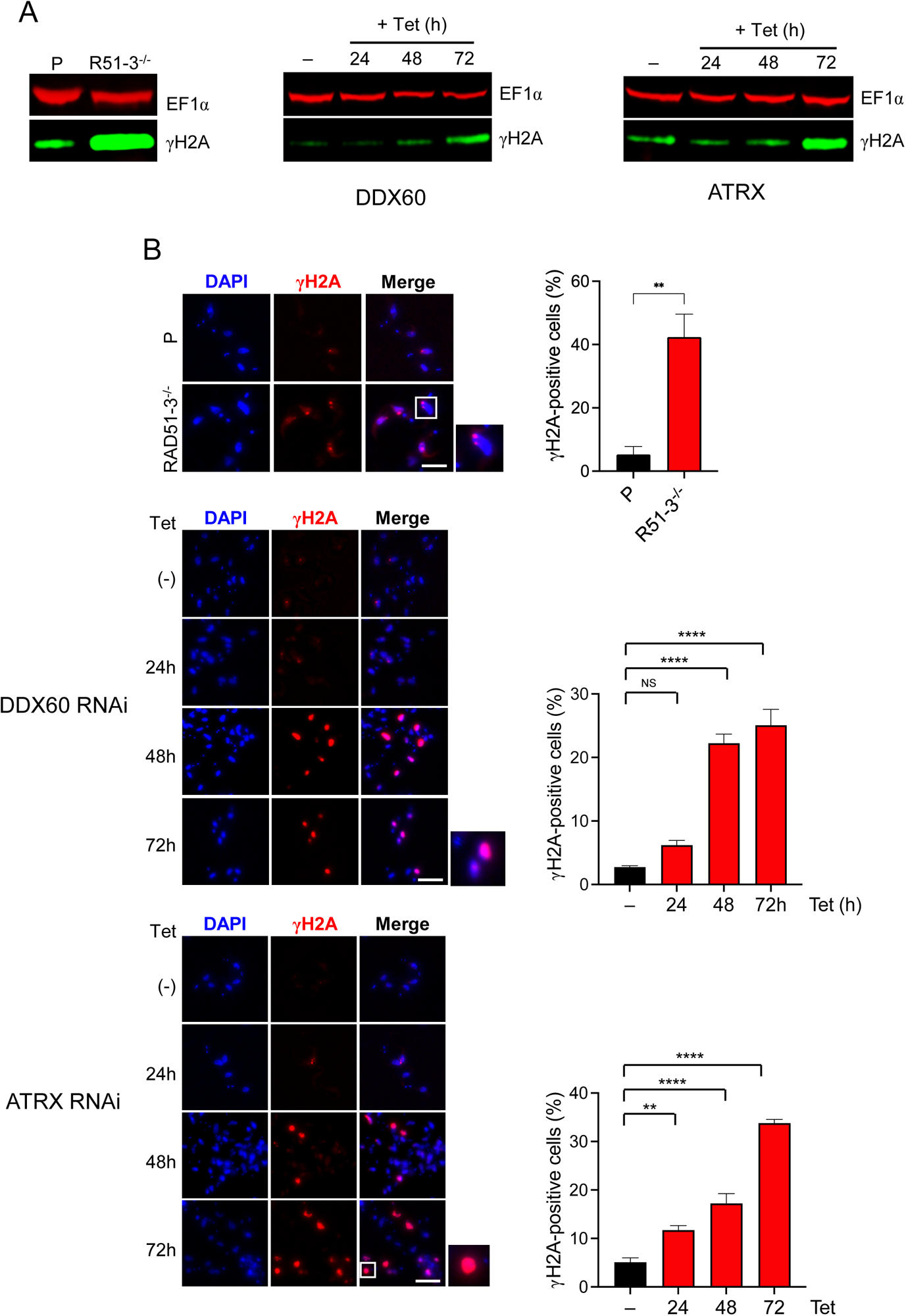
Loss of *T. brucei* RAD51-3, ATRX or DDX60 leads to increased nuclear DNA damage. Anti-γH2A western blot comparing *RAD51-3*-/- mutants with parental (P) cells, and before (Tet-) and after *DDX60* or *ATRX* RNAi induction (Tet+) for 24, 48, and 72 hours (anti-EF1α was used as a loading control). (B) Representative microscopy images of γH2A immunofluorescence in *RAD51-3*-/- mutants and parental (P) cells, and before (Tet-) and after *DDX60* or *ATRX* RNAi induction (Tet+) for 24, 48, and 72 hours. yH2A localisation is quantified in the adjacent graphs, which show the percent of cells with nuclear anti-γH2A signal. Error bars signify SEM from three independent experiments, counting at least 50 cells; statistical significance was determined for the *RAD51-3*-/- mutants using a t-test (**p < 0.01), while statistical significance following RNAi was determined through one-way ANOVA followed by Šídák’s multiple comparisons tests (ns, not significant, **p<0.01, ****p<0.0001).

### Loss of RAD51-3, DDX60 or ATRX alters RNA-DNA hybrid homeostasis

To ask if loss of the putative interactors affects RNA-DNA hybrid dynamics, we performed immunofluorescence with the S9.6. antibody (Figs. 5 and S4). Unlike in mammalian cells (Skourti-Stathaki, 2022; Smolka et al., 2021), the majority of anti-S9.6 signal detected in parental or uninduced RNAi *T. brucei* BSF cells was nuclear (Fig. S4). In addition, and notwithstanding concerns about its effectiveness (Smolka et al., 2021), treatment with *E. coli* RNase H1 significantly reduced S9.6 nuclear fluorescence intensity in the same cells (Figs. 5 and S4), indicating much of the signal represents RNA-DNA hybrids, including R-loops.

**Figure 5.**
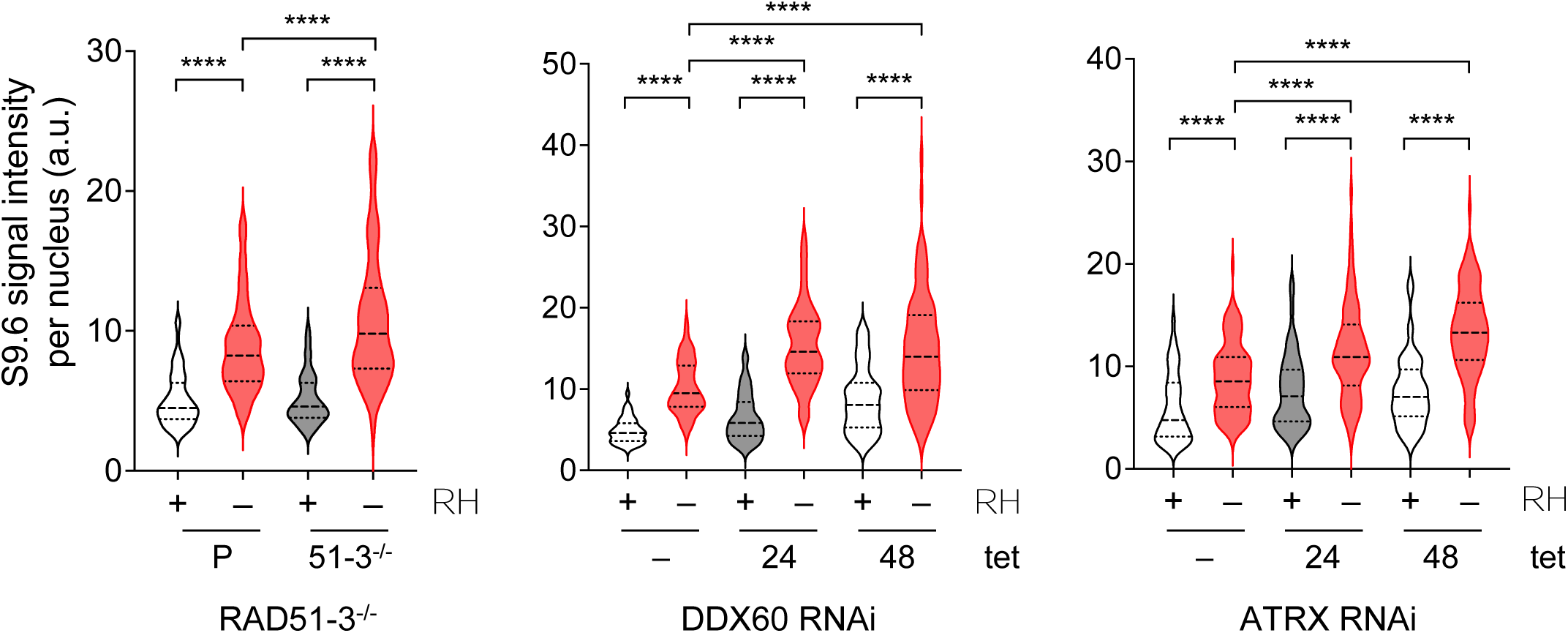
Loss of *T. brucei* RAD51-3, ATRX or DDX60 leads to increased levels of nuclear RNA-DNA hybrids. Violin plots show the intensity of nuclear S9.6 immunofluorescence signal in parental (P) and RAD51-3-/- cells, or before (Tet-) and after *DDX60* or *ATRX* RNAi induction (Tet+) for 24 or 48 hours; in all cases the intensity was measured with (+) and without (-) treatment with *E. coli* RNase H1 (RH). Data is shown in each case for > 100 cells, the median is shown by a heavily dotted line, and the interquartile range by surrounding lightly dotted lines; statistical significance was determined through one-way ANOVA followed by Šídák’s multiple comparisons test (****p < 0.0001).

*RAD51-3-/-* cells displayed significantly increased S9.6 fluorescence compared with the parental cells (Fig. 5). In addition, tetracycline induction of the *ATRX* or *DDX60* RNAi cells led to an increase in S9.6 nuclear signal compared to the uninduced (Fig. 5). In fact, the temporal changes in S9.6 signal appeared to have parallels with the growth curves (Fig. 3B,C) and yH2A immunofluorescence (Fig. 4B: for *ATRX*, median fluorescence increased from 24-48 h after RNAi, whereas for *DDX60* median fluorescence increased by 24 h and was unchanged 48 h post-induction (Fig. 5). Hence, growth impairment and nuclear DNA damage may follow from increased levels of RNA-DNA hybrids due to the loss of the factors. In addition, the findings reiterate a nuclear function for DDX60. Taken together, these data indicate each of these factors acts in RNA-DNA hybrid homeostasis, consistent with their recovery and identification by DRIP-MS.

### Loss of RAD51-3, DDX60 or ATRX alters VSG expression

Given the above evidence linking RAD51-3, DDX60 and ATRX with homeostasis of nuclear RNA-DNA hybrids and with nuclear DNA damage, we next asked if their loss has an impact on VSG switching. MiTat1.2 BSF cells (the strain used for all experiments here) predominantly express VSG221 (also named VSG2) from BES1 [39,40,42]. When wild type MIT1.2 BSF cells are grown in culture, a small proportion (∼1-3%) of cells switch off expression of VSG221 and activate a distinct VSG [39-41,66,67]. To ask if this stochastic switching frequency is altered by loss of the RNA-DNA hybrid interactors, RT-qPCR was first used to assess RNA levels of *VSG221* and four *VSG*s in normally silent BESs (Fig. 6A). In the *RAD51-3*-/- parasites RT-qPCR indicated increased levels of *VSG221* transcript relative to parental cells, and an associated reduction in all *VSG* transcripts from the mainly silent BESs. RNAi of *DD60X* or *ATRX* had the opposite effect: in both cases less *VSG221* transcript was expressed in the 24 h induced populations relative to uninduced, and four or five of the silent BES *VSG* transcripts increased in abundance. These data suggest that loss of RAD51-3 reduces switching away from *VSG221* towards any of the silent BES-resident *VSG*s tested, while loss of DDX60 or ATRX increases switching away from *VSG221* and increases activation of the silent BES *VSG*s. Notably, in the latter cases, switching alteration was detected prior to the significant accumulation of DNA damage or pronounced growth defects beyond 24 hr of RNAi induction.

**Figure 6.**
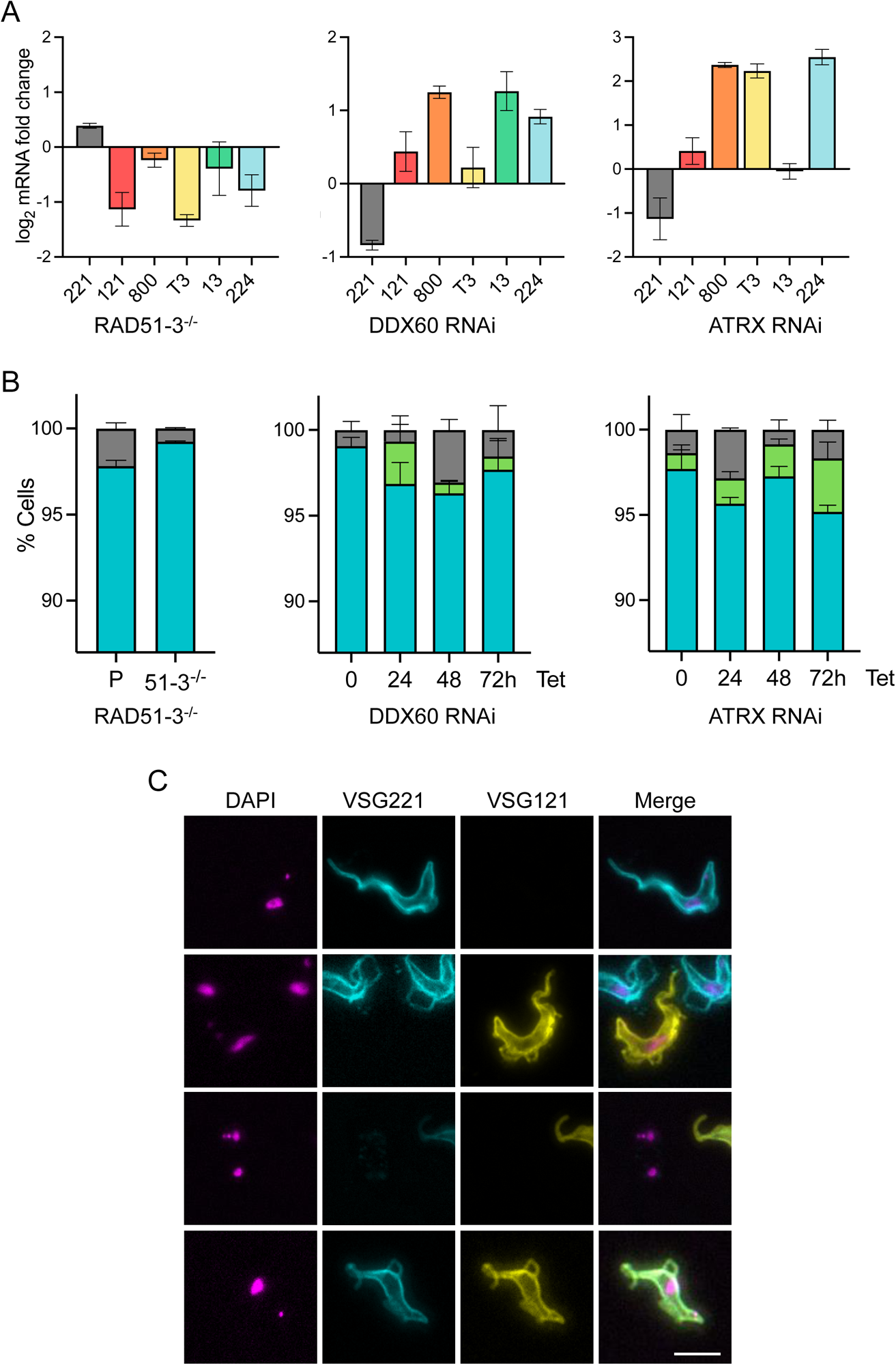
Loss of *T. brucei* RAD51-3, ATRX or DDX60 leads to altered VSG expression. (A) Quantification of the relative levels of *VSG* RNAs, comparing parental and *RAD51-3*-/- mutants, or in cells 24 hours post-induction of RNAi against *DDX60* or *ATRX* compared with uninduced cells; error bars indicate SEM of two independent experiments. (B) Graphical representation of VSG immunofluorescence analysis that shows the proportion of cells staining positive for VSG221, VSG121, both VSGs, or neither VSG in parental (P) and *RAD51-3*-/- cells, or 0, 24, 48 and 72 hrs after RNAi induction (Tet) against *DDX60* or *ATRX*; error bars represent SEM from three independent experiments, counting >300 cells. Sample VSG immunofluorescence images are shown 24 hrs after *ATRX* RNAi; scale bar = 10 µm.

To test the effects on *VSG* expression further, we performed immunofluorescence on live cells with antiserum against VSG221 and VSG121, which is expressed from the predominantly silent BES3 (Fig. 6B). Consistent with the RT-qPCR, more *RAD51-3*-/- cells that express VSG221 were detected in the population compared to the parental line. In addition, while VSG121 was not detected in parental or mutant populations, a reduced number of *RAD51-3*-/- cells were found that did not express either coat protein. These findings are consistent with previous reports that measured VSG switching using *in vivo* immune selection against VSG221 (Dobson et al., 2011; Proudfoot and McCulloch, 2005), and confirm that RAD51-3 loss reduces the efficiency of *T. brucei* VSG switching. VSG221 and VSG121 immunofluorescence provided a fuller explanation of the RT-qPCR analysis after *DDX60* or *ATRX* RNAi (Fig. 6B). Here, the analysis was conducted over 72 hr after RNAi induction and, in both cases, cells were detected that expressed both VSG221 and VSG121 as early as 24 hr (an example is shown in Fig. 6B). Before induction of *DDX60* RNAi no such cells were detected, whereas a small number were seen prior to induction of *ATRX* RNAi and their numbers increased after induction. Taken together, these findings indicate that loss of either factor impairs the gene expression controls that normally operate to ensure only a single *VSG* BES is transcribed, or their RNAi delays the process of transition from expressing VSG221 to VSG121 (and potentially to any new VSG), as seen after loss of DOT1B (Figueiredo et al., 2008). The same effects are seen after loss of RNase H1 or RNase H2A (Briggs et al., 2019; Briggs et al., 2018a), where R-loop levels increase in the *VSG* BESs, indicating a potential link in terms of R-loop homeostasis.

## Conclusions

RNA-DNA hybrids are ubiquitous epigenetic features of all DNA genomes, where their list of functions continues to expand. Understanding this range of functions can be aided by describing the proteins that interact with RNA-DNA hybrids, though such studies have to date only been conducted in mammals. Here, we describe a large cohort of putative RNA-DNA hybrid interactors in the protozoan parasite, *Trypanosoma brucei*, where R-loops have been mapped genome-wide and implicated in both conserved and lineage-specific activities (Briggs et al., 2018b). Consistent with predictions of conserved activities, we find overlap between our data and mammalian studies (Cristini et al., 2018; Kumar et al., 2022), including ribosome-associated proteins and predicted RNA helicases. In fact, our data provide a potentially novel link between these activities, in that we describe a predominantly cytoplasmic DDX (DDX60) that has been found to associate with the ribosome in *T. brucei*’s relative, *T. cruzi* (Bochler et al., 2020), and influences RNA-DNA hybrid levels in the parasite and, moreover, appears to have moonlighting activities in nuclear DNA repair. Our data also provides a number of predicted RNA-DNA hybrid interactors with activities that may reflect particular features of trypanosome gene expression and chromosome biology, including histone variants (Siegel et al., 2009) and kinetochore components (Akiyoshi and Gull, 2014), though it should be noted that tests are needed to determine that these factors do indeed bind RNA-DNA hybrids.

A crucial activity used by *T. brucei* to survive in its mammalian host is antigenic variation, which is driven by transcriptional controls and recombination of *VSG* genes (Faria et al., 2022). Amongst the proteins recovered by DRIP-MS, we have found four that act in VSG switching. Two are key determinants of DNA repair by homologous recombination: RAD51 (Girasol et al, BioRXiv) and RAD51-3. Though both proteins have previously been shown to act in VSG switching by recombination (McCulloch et al., 2015), their interaction with RNA-DNA hybrids provides new mechanistic understanding and builds upon recent work that has implicated R-loops in the reaction (Briggs et al., 2019; Briggs et al., 2018a; Saha et al., 2021; Saha et al., 2020). Two other factors, DDX60 and ATRX, have never before been implicated in VSG switching. How these factors act in antigenic variation is not yet clear, but the observation that their loss, like that of RAD51-3, leads to increased RNA-DNA hybrids levels, while they have distinct effects on VSG switching to the RAD51 paralogue, illustrates that emerging links between R-loops and VSG transcription or recombination deserve further analysis. For instance, the increased levels of RNA-DNA hybrids seen in *RAD51-3* mutants perhaps indicates DNA repair functions that operate more widely than localised activities dedicated to *VSG* recombination.

## Methods

### Trypanosome culture and genetic editing

*Trypanosoma brucei* MiTat1.2 (Lister 427) bloodstream forms were cultured in HMI-9 medium (Life Technologies) supplemented with 10% foetal calf serum (FCS) at 37 °C, 5% CO2, and Lister 427 procyclic forms were cultured in SDM-79 medium with 10% FCS and 0.2% hemin at 27 °C. Bloodstream forms capable of CRSIPR-Cas9 modification were generated by transfection with plasmid pJ1399 (Rojas et al., 2019), containing T7 polymerase and Cas9. Expression of Cas9 was confirmed by RT-PCR (Fig. S4). For epitope tagging and allele knockout, primers were designed with LeishGEdit grown (http://www.leishgedit.net/) and donor sequences were PCR amplified from pPOT plasmid (containing mNeonGreen and blasticidin resistance gene) as previously described (Beneke et al., 2017), along with single guide RNA. Mutant lines were generated by Amaxa transfection with ethanol-purified PCR products and drug-section to generate clonal line, which were confirmed by PCR. Tagged proteins were confirmed by western blot using anti-mNeonGreen antibody (1:1000) and anti-EF1alpha (1:25,000) as a loading control. For inducible RNAi, the BSF Lister 427 derivative 2T1 (Alsford et al., 2005) was used. Target sequences and primers were designed with RNAit (Redmond et al., 2003). PCR amplified fragments were cloned into the pRPaiSL vector plasmid (Alsford and Horn, 2007), which was linearised and transfected to generate RNAi clones, confirmed by RT-PCR.

### RT-qPCR

RNA was extract from 8 x 10^6^ cells using the RNeasy Mini Kit (Qiagen) protocol. Genomic DNA was digested on-column for 15 min at room temperature using the RNase-free DNase I Set (Qiagen). First-strand complementary DNA (cDNA) synthesis cDNA was generated from 500 ng total RNA using SuperScript™ IV First-Strand Synthesis System (Invitrogen), following the manufacturer’s protocol using random hexamer primers. Previously used primer sequences were used for VSG RT-qPCR (Tiengwe et al., 2012). Each primer pair target was run in two biological replicates and three technical replicates for each cell line. A 20 μL reaction contains 1X SYBR. Green PCR Master Mix (Applied Biosystems), 250 nM of forward and reverse primers and 1 μL cDNA. All qPCR experiments were run in 7500 Real-Time PCR system (Applied Biosystems) using the following cycling conditions: 1 cycle at 95 °C for 10 min, followed by 40 cycles of 95 °C for 15 sec and 60 °C for 1 min. Fluorescence intensity was measured at the end of each extension step (60 °C for 1 min). For normalization across different samples, actin amplification was used as endogenous control. For calculation of relative mRNA levels, the 2-ΔΔCt method was used (Livak and Schmittgen, 2001).

### Fluorescence microscopy

For imaging mNeonGreen fluorescent proteins, ∼8 x 10^6^ parasites were pelleted by centrifugation at 800 rcf for 7 min and resuspended in FCS-free HMI-9 media with 1 µg/mL Hoechst 33342 (Sigma-Aldrich). Cells were pelleted again, resuspended in of 0.05% (v/v) formaldehyde in FCS-free HMI-9 to immobilise parasite flagella and adhered to a Poly-L-Lysine coated slide for immediate imaging. For immunofluorescence analysis of DNA damage, washed parasites were fixed in 4% formaldehyde in vPBS for 10 min and permeabilised with 0.1% IGEPAL CA-630 for 10 min. Cells were washed in vPBS and adhered to a slide, before incubating in PSB + 1% (w/v) glycine 5 min and blocked with 1% bovine serum albumin (BSA) for 1 hr. Staining was performed with α-yH2A primary (1:1,000) and α-rabbit Alexa Fluor® Plus 488 (1:1,000) secondary antibodies diluted in 1% BSA. Slides were washed 1X PBS before mounting with 5 µL DAPI Fluoromount-G® (Southern Biotech) and imaging. For imaging DNA-RNA hybrids, parasites were instead fixed in 70% ice-cold methanol for 1 hr, permeabilised with 0.5% v/v Triton X-100 for 10 min and blocked with 1X PBS, 0.01% v/v Tween-20, 0.1% w/v BSA for 1 hr at 37°C. S9.6 (Kerafast) primary (1:1,000) and Alexa Fluor Plus 488 Goat anti-Mouse IgG (H+L) (ThermoFisher) secondary (1:3,000) antibodies were diluted in blocking solution for staining in suspension while shaking before adhering to slides. VSG immunofluorescence analysis was performed following the protocol of Glover et al. (Glover et al., 2016) Briefly, formaldehyde fixed parasites were adhered to glass slides and blocked with 50% FCS in PBS for 45 min before staining with primary anti-VSG (1:10,000) and secondary Alexa Fluor (1:1,000) antibodies and mounting with DAPI Fluoromount-G® (Southern Biotech). Imaging was performed with an Axioscope 2 widefield fluorescence microscope (Zeiss) using a 63x/1.40 oil objective, or a Leica DiM8 widefield fluorescence microscope to acquire Z-stacks of 5 µm thickness in 25 sections. To quantify fluorescence intensity, images were obtained using the same exposure times and were later processed on Fiji/ImageJ (Schindelin et al., 2012) using the same parameters (http://imagej.net/Rolling_Ball_Background_Subtraction).

### DRIP-MS

DRIP-MS approach was adapted from Cristini et al (Cristini et al., 2018). 2.5 x 10^8^ mid-log growth phase parasites were pelleted and washed in 1 ml of 1X PBS, before incubation in 1 mL of lysis buffer (80 mM KCl, 5 mM PIPES pH 8.0, 0.5% Nonidet P-40 substitute) on ice for 20 min. The pellet was homogenized using a tight-fitting dounce homogenizer until the nuclei were release, as checked with microscope. Nuclei were pelleted (2,400 rcf for 10 min, 4 °C) and resuspended in 200 μL of RSB sonication buffer [1X RSB buffer (10 mM Tris-HCl pH 7.5, 200 mM NaCl, 2.5 mM MgCl_2_), 0.2% sodium deoxycholate, 0.1% SDS, 0.05% sodium lauroyl sarcosinate, 0.5% Triton X-100] before sonicating for 10 min in the Diagenode Bioruptor using cycles of 30 sec ON and 90 sec OFF. The sonicated nuclei were then diluted with 600 μL RSB+T buffer (1X RSB buffer, 0.5% Triton X-100) plus 0.8 ng RNase A to degrade single-stranded RNA. For benzonase control IPs, 1U/μL benzonase was added and incubated for 30 min at 37°C. 20 μL was reserved as input control samples. Pre-prepared Protein A Dynabeads (incubated with 5 μg of S9.6 antibody (Millipore) in 1X PBS plus 0.5% BSA overnight at 4°C), were washed in 1X PBS + 0.5% BSA and suspended in the sonicated sample. Samples were incubated on a rotor at 4°C for 2 hrs to bind DNA-RNA hybrids and interacting proteins. Beads were then wash in 1 mL cold RSB+T buffer (4 times) and 1 mL cold RSB buffer (twice), before collecting by centrifugation at 1,000 rcf for 3 min at 4°C. Beads were retained using a magnetic rack and supernatant was discarded. Beads were suspended in 15 μL loading buffer (1X NuPAGE LDS loading buffer, 1X PBS, 0.1% β-mercaptoethanol) for 10 min at 70°C. Beads were pelleted and supernatant, containing eluted proteins collected.

Proteins were resolved using sodium dodecyl sulfate-polyacrylamide gel electrophoresis (SDS-PAGE) and stained using InstantBlue® Coomassie Protein Stain (Abcam). Gel segments only found in S9.6 IP samples and not benzonase controls were excised for Nanoflow HPLC Electrospray Tandem Mass Spectrometry (nLC-ESIMS/MS), performed by Proteomics Facility of Glasgow Polyomics. Briefly, the gel pieces were de-stained by incubation in 500 μL of 50% acetonitrile in 100 mM ammonium bicarbonate for 30 min on a shaker, dehydrated by incubation in acetonitrile for 10 min then dried a vacuum centrifuge. Trypsin was added to rehydrate the gel pieces and digested overnight at 37°C, before gel pieces were pelleted. The supernatant containing peptides was dried in a vacuum centrifuge and solubilized in 20 μL 5% acetonitrile with 0.5% formic acid.

Trypsinized peptide samples were analyzed using nanoscale liquid chromatography coupled to electrospray ionization tandem mass spectrometry (nLC-ESI–MS/MS). Online detection of peptide ion was by electrospray ionization mass spectrometry using an Orbitrap Elite MS (Thermo Scientific). Peptides were separated on a PepmapTM C18 reversed phase column (3 μm, 100 Å, 75 μm × 50 cm) (Thermo Scientific). Samples were fractionated with mobile phase A consisting of 0.1% (v/v) formic acid in water and mobile phase B consisting of acetonitrile (80% v/v) and water (20% v/v). The peptide separation was performed at a fixed solvent flow rate of 0.3 µL/min, using a gradient of 4–100% mobile phase B over 120 min. The Orbitrap EliteTM MS acquired full-scan spectra in the mass range of m/z 300–2000 Da for a high-resolution precursor scan at a set mass resolving power of 60,000 (at 400 m/z). Collision-induced dissociation was performed in the linear ion trap with the 20 most abundant precursors using rapid scan mode.

Proteins were identified using the Mascot search engine (v2.6.2, Matrix Science) by interrogating MS data against protein databases of *Trypanosoma brucei* Lister 427. A mass tolerance of 10 ppm was allowed for the precursor and 0.3 Da for MS/MS matching, with the following search parameters: trypsin enzyme specificity, allowing one missed cleavage; cysteine carbamidomethylation was selected as fixed modification, while N-terminal carbamidomethylation, asparagine-glutamine deamidation, tyrosine iodination, and methionine oxidation were selected as variable modifications. Proteins within a significance score of p < 0.05 and with at least one unique peptide were considered. Four and two independent biological replicates were generated for BSF and PCF *T. brucei,* respectively. Corresponding emPAI values were compared between IP and benzonase controls; proteins with mean log2 fold-change >0 across all biological replicates constituted the RNA/DNA hybrid interactome for each form. BSF-specific interactome was defined as proteins with > 0 log2 fold-change in BSF vs PCF.

### Homology searches, domain analysis, and phylogenetics

To look for putative homologs of Tb927.3.2600 and Tb927.3.5440 in Discoba and elsewhere, sequence- and HMM-based similarity searches were done in HMMER (Potter et al., 2018) and HHpred (Steinegger et al., 2019; Zimmermann et al., 2018). KEGG Orthology (KO) scores, which guided our searches for homologs and phylogenetic analyses, were obtained with KofamKOALA (Aramaki et al., 2020). To determine the Pfam domain organisation of every analysed protein, sequences were run through InterProScan (Paysan-Lafosse et al., 2023). Helicase ‘core’ regions were defined according to Fairman-Williams et al. (Fairman-Williams et al., 2010). The regions of sequence similarity shown in Fig. 2, were obtained from HHpred searches using Tb927.3.2600 and Tb927.3.5440 as queries against HMM databases representing the proteomes of *D. melanogaster* and/or *H. sapiens*; for every protein pair that we considered, we defined their ‘region of similarity’ as that corresponding to their whole alignment as retrieved by HHpred. For the phylogenetic analysis of Tb927.3.2600, we first collected putative Tb927.3.2600 homologs from across Discoba (searches were done in publicly available genomic and transcriptomic datasets), as well as representative orthologs of Tb927.3.2600’s top 4 hits from KofamKOALA, namely DDX60 (KO entry: K20103), MTR4 (K12598), ISE2 (K26077), and HelY (K03727), all of which are members of the Ski2-like family of SF2 helicases(Bobik et al., 2017; Fairman-Williams et al., 2010; Uson et al., 2015). We then used MAFFT-DASH L-INS-i (Katoh and Standley, 2013; Rozewicki et al., 2019) to align each of the 5 groups of protein sequences (Tb927.3.2600 and its putative homologs in Discoba; DDX60; MTR4; ISE2; and HelY) separately; in each case, alignment was followed by manual trimming in Jalview (Waterhouse et al., 2009), down to the helicase ‘core’ plus short flanking regions (approximately 50 to 100 aa, depending on sequence conservation within each group). All resulting subsequences were then aligned with MAFFT-DASH L-INS-i, and the obtained alignment trimmed with trimAl (Capella-Gutierrez et al., 2009) using a column gap threshold of 10% (‘gt - 0.1’ option). This yielded a ‘final alignment’, which was used as input in an IQ-TREE (Minh et al., 2020) run to infer a maximum-likelihood phylogenetic tree; the best-fit evolutionary model was selected by ModelFinder (in addition to default ‘standard’ models, several mixture models, e.g. EX_EHO, were included in ModelFinder’s search), and both SH-aLRT (Guindon et al., 2010) and UFBoot2 (Hoang et al., 2018) were performed with 1000 replicates each. For the phylogenetic analysis of Tb927.3.5440, a similar protocol was followed. Tb927.3.5440’s top 4 hits from KofamKOALA were ATRX (K10779), ARIP4 (K10876), RAD54 (K10875) and RAD54B (K10877), all of which are members of the Rad54-like grouping within the Snf2 family of SF2 helicases (Flaus et al., 2006). KO’s RAD54 entry confusingly includes a mixture of RAD54 and DRD1 orthologs, which we were able to identify in preliminary trees; because DRD1 is also part of the Rad54-like grouping, we included it in our analysis. Phylogenetic trees were rooted with MAD (Tria et al., 2017), and visualized and edited in iTOL (Letunic and Bork, 2021).

### Data availability

DRIP-mass spectrometry proteomics data have been deposited to the ProteomeXchange Consortium via the PRIDE partner repository, with the dataset identifier PXD042146.

## Acknowledgements

This work was funded by a Wellcome Trust Investigator Award (224501/Z/21/Z) to R.M., a Philippine Council for Health Research and Development (DOST-PCHRD) award to M.G., a Welcome Trust Sir Henry Wellcome fellowship (218648/Z/19/Z) to E.M.B., a Wellcome Institutional Strategic Support Fund (ISSF3) award held at the University of Glasgow (204820/Z/16/Z) to E.M.B. and R.M., and by BBSRC awards (BB/N016165/1, BB/R017166/1 and BB/W001101/1) to R.M. and C.A.M. The Wellcome Centre for Integrative Parasitology is funded by a Wellcome Trust Strategic Award (104111/Z/14/Z/A). We thank Suzanne McGill and all staff at Glasgow Polyomics for support in mass spectrometry analysis.

## Supplementary figure legends

**Figure S1.**
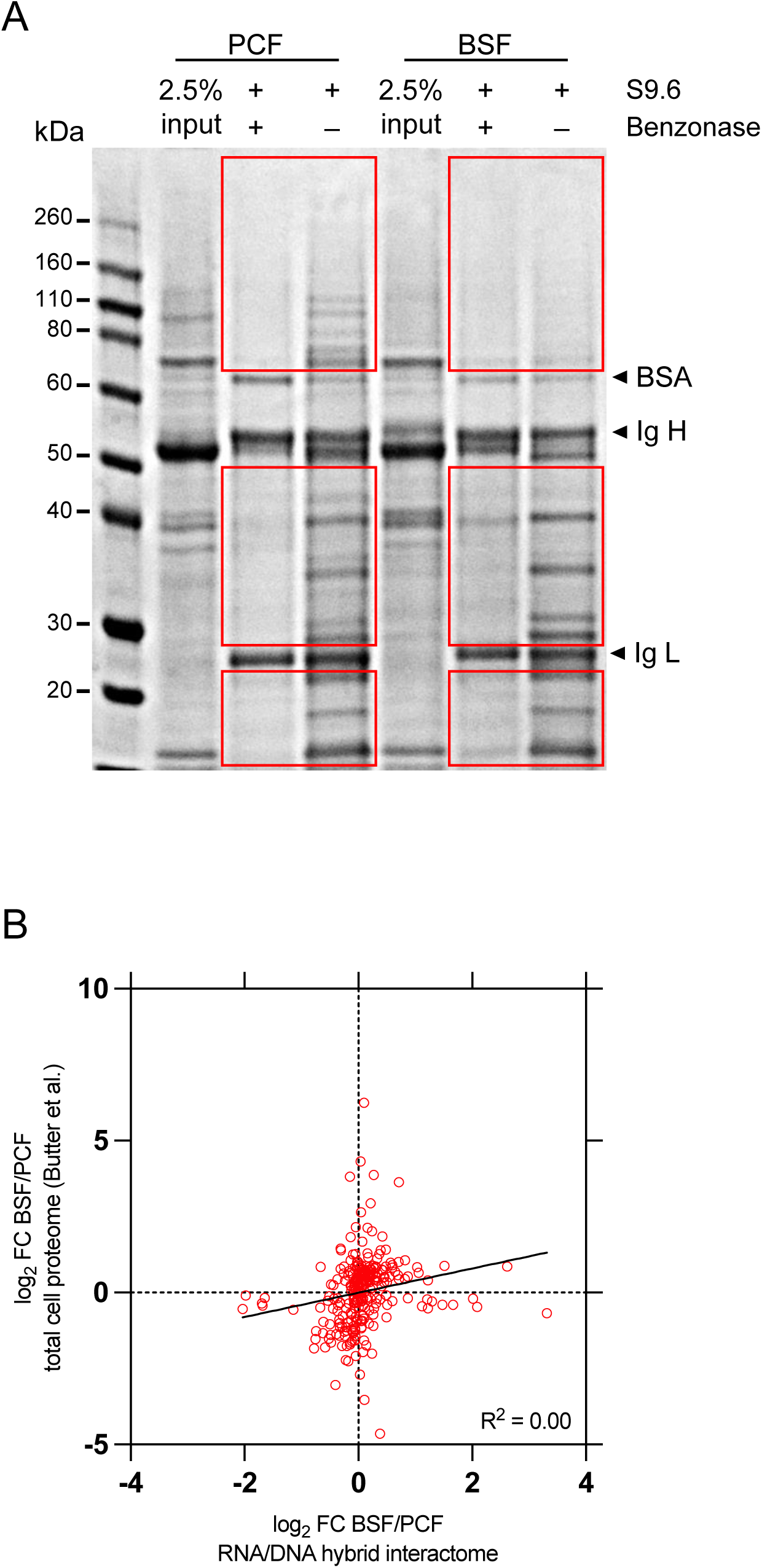
DRIP-MS enriches for a subset of proteins from the total cell proteomes of both BSF and PCF *T. brucei.* (A) Representative Coomassie-stained gel of eluted proteins from DRIP with or without benzonase treatment. BSA indicates Bovine Serum Albumin used to block protein A dynabeads; IgH and IgL indicate heavy chain and light chain of the S9.6 antibody, respectively. Red boxes indicate regions excised for mass spectrometry. (B) Comparison of log2 fold enrichments of proteins in the combined RNA-DNA hybrid interactomes (this study) and the total cell proteome obtained from the dataset in a study by Butter et al (Butter et al., 2013).

**Figure S2.**
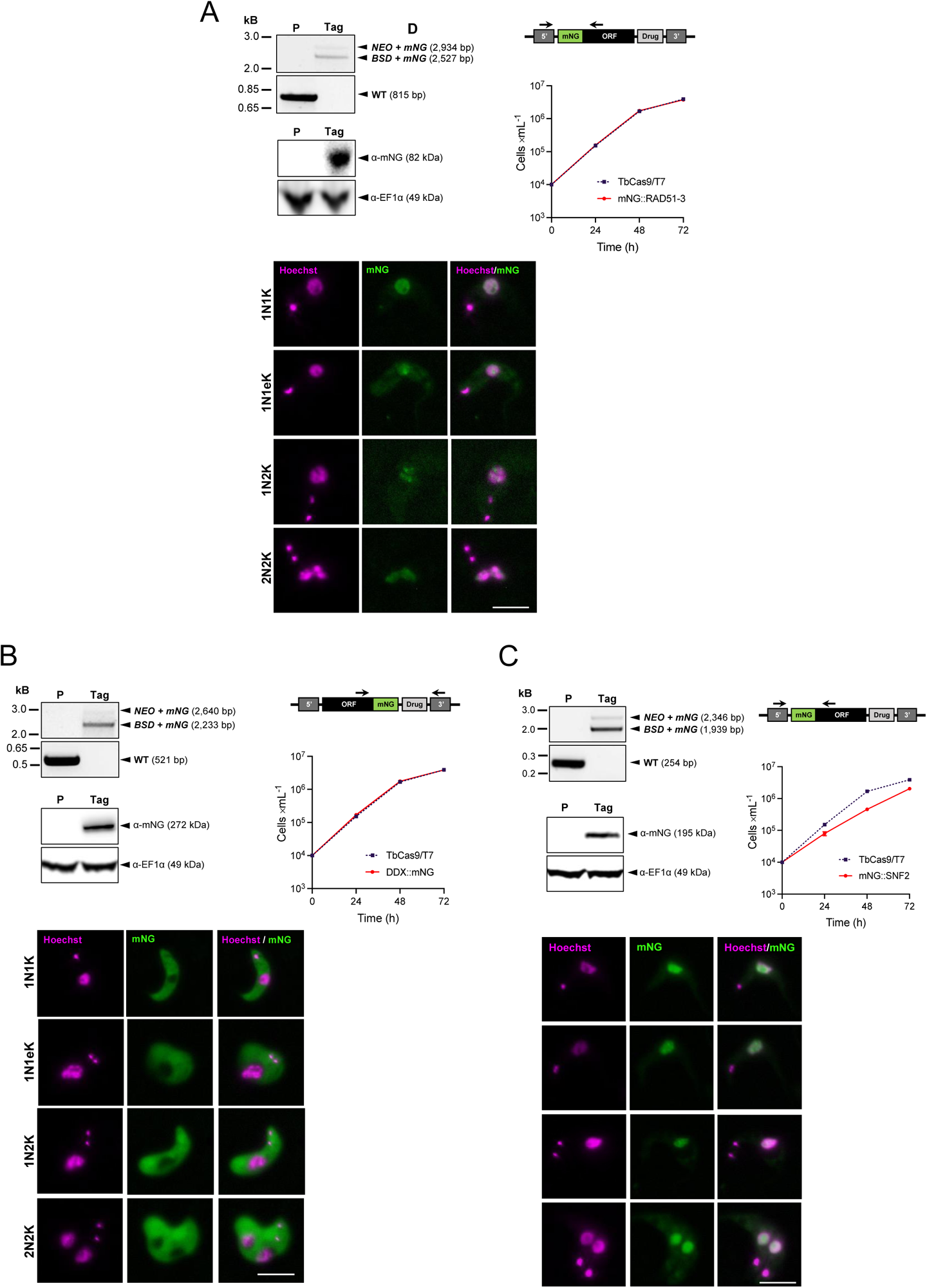
Localisation of *T. brucei* RAD51-3, ATRX and DDX60. For each of RAD51-3 (A), DDX60 (B) and ATRX (C), the following are shown: PCR showing transformants with both alleles tagged with mNeonGreen (mNG; P, parental TbCas9/T7; Tag, mNG transformant; WT, untagged allele); annealing sites of primers used for PCR; western blot analysis of whole-cell protein extracts probed with anti-mNeonGreen antibody, anti-EF1α antibody was used as a loading control (expected sizes of the mNG-tagged proteins are shown); growth analysis of the mNG transformant compared to the parental TbCas9/T7 cell line (error bars represent SEM from three independent experiments); and representative microscopy images of live fluorescence imaging of mNG transformant cells at different cell cycle stages (scale bar = 5 µm).

**Figure S3.**
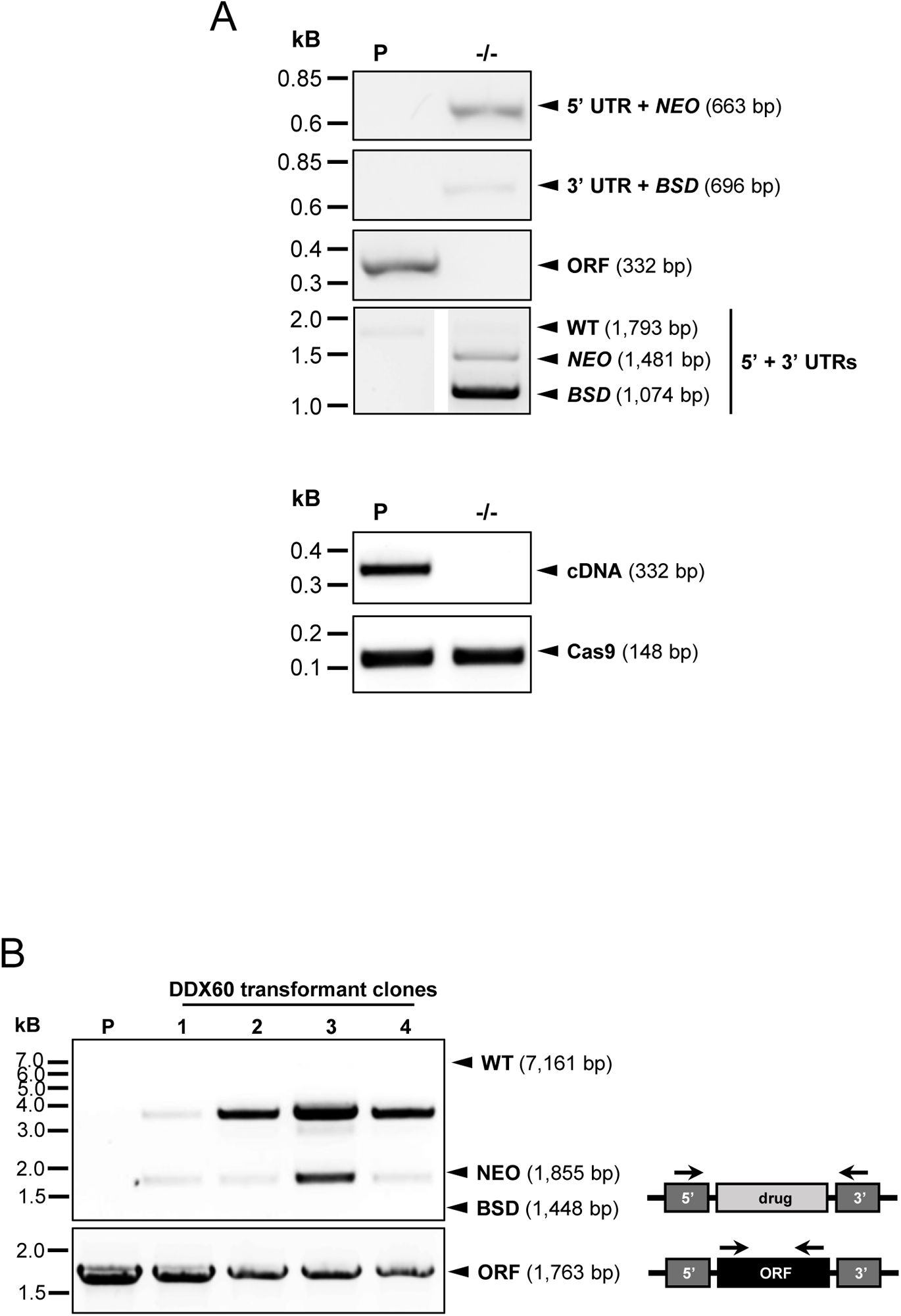
Attempts to make null mutants of *RAD51-3* and *DDX60* by CRISPR-Cas9. (A) PCR of a *RAD51-3-/-* transformant and parental (P) TbCas9/T7 cells, demonstrating the expected integration of *NEO* and *BSD* knockout constructs and the corresponding loss of the *RAD51-3* ORF, as well as RT-PCR results demonstrating the loss of *RAD51-3* RNA (*Cas9* was used as a control). (B) PCR to analyse integration of NEO and BSD knockout constructs in a number of *DDX60* transformant relative to parental TbCas9T7 cells; sizes of PCR products for allele replacement integrations are shown, as well as for the unaltered allele (WT), and approximate annealing sites of primers used for PCR are indicated.

**Figure S4.**
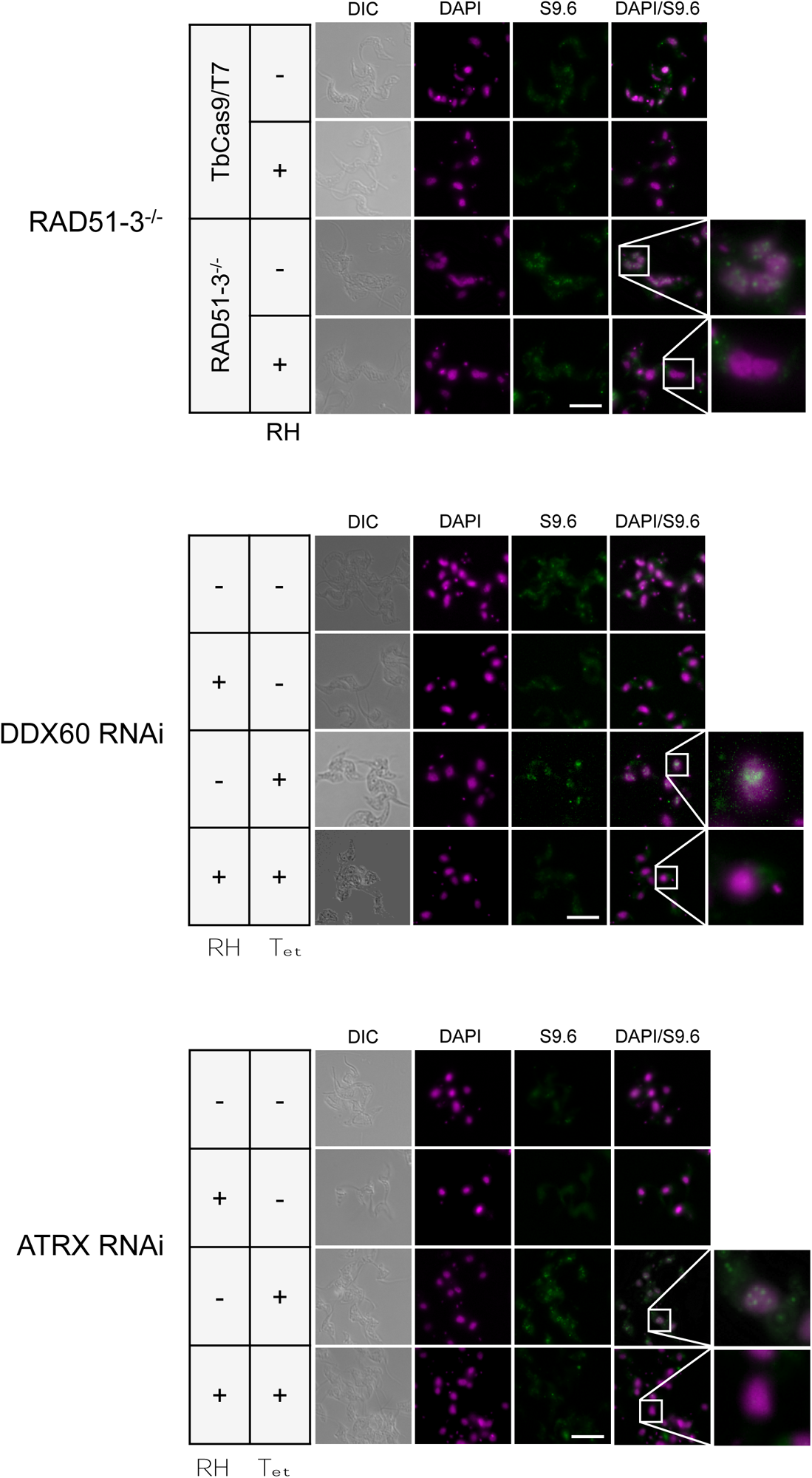
Loss of *T. brucei* RAD51-3, ATRX or DDX60 leads to increased levels of nuclear RNA-DNA hybrids. Representative images of S9.6 immunofluorescence, comparing *RAD51-3-/-* and parental TbCas9T7 cells, and cells grown with (Tet-) or without (Tet+) RNAi induction against *DDX60* or *ATRX*; in all cases, images are shown with or without *E. coli* RNase H1 (RH) treatment. Inset images depict zoomed-in regions showing nuclear distribution of R-loops. Scale bar before zoom = 10 µm.

**Table S1.** Analysis of proteins recovered by DNA-RNA immunoprecipitation and identified by mass spectrometry, showing four bloodstream (BSF) and two procyclic form (PCF) experiments relative to two and one, respectively, benzonase treated controls (ctrl).

